# Understanding trivial challenges of microbial genomics: An assembly example

**DOI:** 10.1101/347625

**Authors:** Delphine Lariviere, Han Mei, Mallory Freeberg, James Taylor, Anton Nekrutenko

## Abstract

The perceived “simplicity” of bacterial genomics (*these genomes are small and easy to assemble*) feeds the decentralized state of the field where computational analysis standards have been slow to evolve. This situation has a historical explanation. In cases of human, mouse, fly, worm and other model organisms there have been large sustained multinational genome sequencing efforts and analysis consortia such as the 1,000 genomes, ENCODE, modENCODE, GTEx and others. These resulted in development and proliferation of common tools, workflows, and data standards. This is not the case in microbiology. After the development of highly parallel sequencing methodologies in mid-2000s bacterial genomes no longer required initiatives of such scale. The flipside of this is the extreme heterogeneity of approaches to many well established microbial genomic analysis problems such as genome assembly. While competition amongst different methods is good, we argue that the quality of data analyses will improve if cutting edge tools are more accessible and microbiologists become more computationally savvy. Here we use genome assembly as an example to highlight current challenges and to provide a possible solution.

---

We develop and maintain a popular genomic analysis platform—Galaxy^1^. In the course of this long running project we became keenly aware of the fact that developing computational platforms and using them are distinct activities: developer cannot accurately anticipate the needs of an experimental user. But how to ensure that software development priorities are in alignment with the needs of real users? Perhaps the best approach is to select a common type of analysis and to understand what is involved in completing one. One of the authors of this manuscript is an experimentalist without prior computational expertise working on in vitro evolution project stemming from our previous work with *E. coli* C and bacteriophage ϕX174^2^. The genome of *E. coli* C has not been previously determined and by sequencing it we wanted to discover, enumerate, and attempt to solve any hurdles that arise. So we set our purely experimental colleague on a journey to perform assembly without any explicit computational help. After this has been painstakingly accomplished we modified Galaxy system to account for all analytical idiosyncrasies discovered during this effort and created a detailed interactive tutorial (https://goo.gl/xP7jyn; our ultimate goal is to create multiple similar tutorials on common types of microbial genomic analyses). The set of issues that we discovered was very illuminating for understanding hurdles preventing microbiologists from embracing an ever growing set of free, community-supported, open-source analysis tools.

The first fully sequenced genome of a free living organism was that of *Haemophilus influenzae* ^3^. It was a result of a large collaborative effort that involved the development of a dedicated genomes assembler ^4^. Since that time numerous advances in genome sequencing and assembly have transformed life sciences. Today there are established experimental protocols for preparing sequencing libraries for a variety of currently available platforms. Similarly there are established, well tested, open-source software tools for assembly of sequencing reads of various lengths and error profiles into complete genomic sequences ^6,7^ As a result bacterial genome sequencing is a common task: for *E. coli* alone over 10,800 genome assemblies (June 2018; both complete and partial) have been deposited to the National Center for Biotechnology Information (NCBI) microbial genome database since the publication of the K-12 sequence in 1997 ^5^. Given all these advances how easy is it to actually assemble a bacterial genome?

A microbiologist experienced in molecular biology techniques will have no trouble isolating genomic DNA and preparing sequencing libraries. To sequence the C-1 strain (Coli Genetic Stock Center #3121) of *E. coli* used in our experimental evolution study we chose two commercially available technologies: Illumina and Oxford Nanopore. Illumina is the current standard in short read high coverage sequencing and is widely accessible through institutional core facilities, commercial sequencing service providers and individual labs. Oxford Nanopore (ONT) is a maturing single molecule sequencing technology that produces long reads with relatively high error rate. Combining high accuracy short Illumina reads with error prone but long ONT reads allows performing hybrid assembly ^9^ yielding accurate and, often, complete bacterial assemblies. We specifically chose ONT over the other more established long read single molecule sequencing approach, Pacific Biosciences (PacBio), because ONT’s perceived appeal to small labs. This appeal is based on low up-front cost and small physical footprint of ONT sequencer—MinION. Additionally, ONT still does not have firmly established data processing and analysis workflows and we were interested in experiencing this first hand. Using Illumina TruSeq library preparation protocol on a MiSeq machine and R7 ONT chemistry on a MinION MK1b device we generated 9,345,897 250 bp Illumina read pairs and 14,093 long nanopore reads with maximum length of 27.5 kb. Both Illumina and ONT data are deposited to Short Read Archive under accession SRP131264 and can also be accessed from Zenodo (DOI:10.5281/zenodo.1257429).

Sobering reality begins immediately upon receiving the data from the MinION machine. While Illumina data comes in well established fastq format ^10^ ONT data poses a challenge. In our case the data was obtained in fast5 format, in which every read is in a separate file. Since our run yielded 14,093 reads this translated into 14,093 files that occupy over 16Gb of disk space. Moving and manipulation of such file collections is inevitably a challenge and to be useful for downstream analyses they need to be converted to fastq format as well. This can be achieved with a free, community-developed package called Poretools ^11^. Poretools can be installed and used by individuals familiar with any flavor of UNIX environment (e.g., MacOS and Linux) but pose challenge for naive users (like one of the co-authors of this manuscript) and owners of Windows PCs. To address the need to manipulate ONT data we have wrapped Poretools for Galaxy and used them to generate fastq representation of nanopore reads. We retained 12,738 high quality 2D reads ranging from 1 to 27.5kb in length (*N*50 = 8,808). Illumina reads were of sufficiently high quality and did not require any additional processing.

Next, one needs to decide which method to be used for read assembly. One possible way to obtain the information necessary to make this decision is to see which tools were used to assemble existing genomes. At the time of writing NCBI microbial genome database contained 10,596 genomes labelled as “complete” (June 2018). Over half of these (5,849) have been deposited to NCBI in the past three years (after January 2017). Of these 5,404 contained useful metadata with 107 genomes listing both “Illumina” and “Oxford Nanopore” as sequencing technologies used. The majority (74) used Unicycler ^8^ with SPAdes ^12^ being second most widely used assembly tool. In reality however, this simple numerical analysis we just described cannot be easily conducted by most experimentalists. It requires downloading tabular datasets from NCBI, writing scripts that would parse these data and download files containing assembly information, processing these files, and generating the report. As a result most of our experimental colleagues will resort to random clicking with, statistically speaking, chances of finding one of 74 genomes assembled with Unicycler among 10,596 being slim. But for the sake of continuing our experiment let us settle on using Unicycler for assembly. Unicycler uses SPAdes as the key component of its assembly process. It is a are pre-configured assembly pipeline that combines read error correction (error correction), various assembly tools tuned for different types of sequencing data such as short accurate reads and/or long noisy reads, and post-assembly steps such as polishing, variant calling, and assembly rotation into a single pipeline.

Now that we have the data and know which assembly tool to use it should be trivial to perform the assembly. It turns out not be the case: our modestly sized dataset (just over 9 million read pairs) cannot be assembled on an ordinary lab desktop—assembly is both memory and CPU intensive and while bacterial genomes have the advantage of being compact, these issues still persist. There is a free solution to this challenge. The United States operates a collection of large NSF-funded high performance computing systems dedicated to scientific computing—https://www.xsede.org/. Free CPU-time and disk space allocations can be obtained on XSEDE resources by writing straightforward, rapidly reviewed applications. However, these resources are underused by experimental life scientists as their proper utilization requires familiarity with scientific computing practices. Galaxy is designed to be able to distribute analyses across geographically distributed heterogeneous computational resources such as various XSEDE components. Galaxy’s Unicycler analyses run on the Bridges high performance computing resource at the Pittsburgh Supercomputing Center—a large shared-memory XSEDE resource ideally suited for genome assembly (https://www.psc.edu/bridges). Using Bridges Unicycler assembled Illumina and ONT sequencing reads into two contigs 4,576,290 and 5,386 bases in length, respectively (assembly took approximately 4 hours utilizing 80 CPUs). The larger contig represents the complete genome of *E. coli* C-1, while the smaller is 100% identical to the sequence of bacteriophage *φ*X174 used as a spike-in in Illumina sequencing protocols. To assess the quality of the new assembly and to annotate genes we integrated into Galaxy and used Quast ^13^ and Prokka ^14^ tools, respectively.

The newly created assembly should now be analyzed in the comparative context: are there large insertions or deletions that differentiate our strain from those already sequenced? Such an analysis involves alignment of our assembly against already sequenced genomes. Thus the initial logistical challenge is locating and obtaining these already sequenced genomes. NCBI provides means for doing this. For example, at the time of writing there were 592 complete *E. coli* genomes (https://www.ncbi.nlm.nih.gov/genome/genomes/167). NCBI allows users to download a table with information about individual assemblies. This table contains web addresses pointing to a remote disk folders containing genomic sequences and annotations. But these web addresses are partial (do not contain names of the files) and to download DNA sequence files each address needs to be modified. This will have to be done 592 times. Such editing can potentially be performed in a spreadsheet application, but this still does not make easy to download all the files automatically. To address this we designed “rule-based” uploader allowing users to fetch complex collections of multiple files (https://vimeo.com/271328293). Using this tool we uploaded all 592 genomes into Galaxy.

Similarly to assembly problem large scale analyses of sequencing data require robust computational infrastructure and, again, infrastructure provided by XSEDE is superbly appropriate. At this point we needed to align our assembly against 592 finished *E. coli* genomes. Alignment is computationally distinct from assembly in that it does not require large memory allocation and can be performed on conventional clusters. For that purpose Galaxy takes advantage of another high performance shared XSEDE resource—JetStream (https://jetstream-cloud.org/). As is the case for assembly there is a number of established tools for performing alignments with BLAST ^15^ being the most well known. In the case of this analysis BLAST has to be downloaded and installed, a BLAST database has to be created from the 592 genome set and searched against. None of these steps are particularly difficult given familiarity with UNIX environment but are immensely challenging otherwise. We used a different, BLAST-like aligner LASTZ (https://lastz.github.io/lastz/) designed for long genomic sequences by simultaneously starting 592 jobs in Galaxy each running for approximately 5 min. All alignment data were then combined into a single dataset.

Up to this point, most difficulties experienced by a computational novice were related to choosing appropriate tools, finding sufficiently powerful infrastructure, and downloading bulk data collections from NCBI. What comes next is different. Manipulations we are about to describe are straightforward, but they are punctuated by numerous, ridiculously unsophisticated, issues. These trivial challenges are chiefly responsible for the perception that data analysis in biology is unpleasant and difficult.

Alignment dataset that we just produced represents coordinates of regions that are highly similar between our assembly and each of the 592 *E. coli* genomes. This is a very straightforward dataset to deal with in all aspects except one—size—it contains ~13,000,000 rows. This is too much for Excel or Google Sheets to process. The majority of rows in this file are likely to be spurious alignment hits with low similarity, which need to be filtered. But how to assess what should be filtered out from a dataset so large? For someone familiar with scientific computing sqlite or Pandas ^16^ will offer a solution. Alternatively, Galaxy does not have restriction on file size. To identify erroneous alignments one can plot the relationship between alignment length and its identity. 13 million rows is still too much data to plot. Instead we generated a random subset with only 10,000 rows and plotted the length/identity relationship (Fig. 1A). The majority of alignments are represented by spurious hits below 10,000 bp and 90% identity and can be removed. This retains only 0.43% of the original data and creates a dataset that is much easier to manipulate. In particular we can aggregate data: for each of the 592 *E. coli* genomes, we compute the number of contiguous alignments, their total length, and plot them against each other (Fig. 1B). Such aggregation can be quickly performed with tools like Datamash (https://www.gnu.org/software/datamash/) in Galaxy. This identified a cluster of three genomes, LT906474.1 (strain NCTC122), CP024090.1 (strain WG5), and CP020543.1 (another instance of *E. coli* C), that have high total alignment length produced by just a handful of alignments (Fig. 1B).

**Figure 1.**
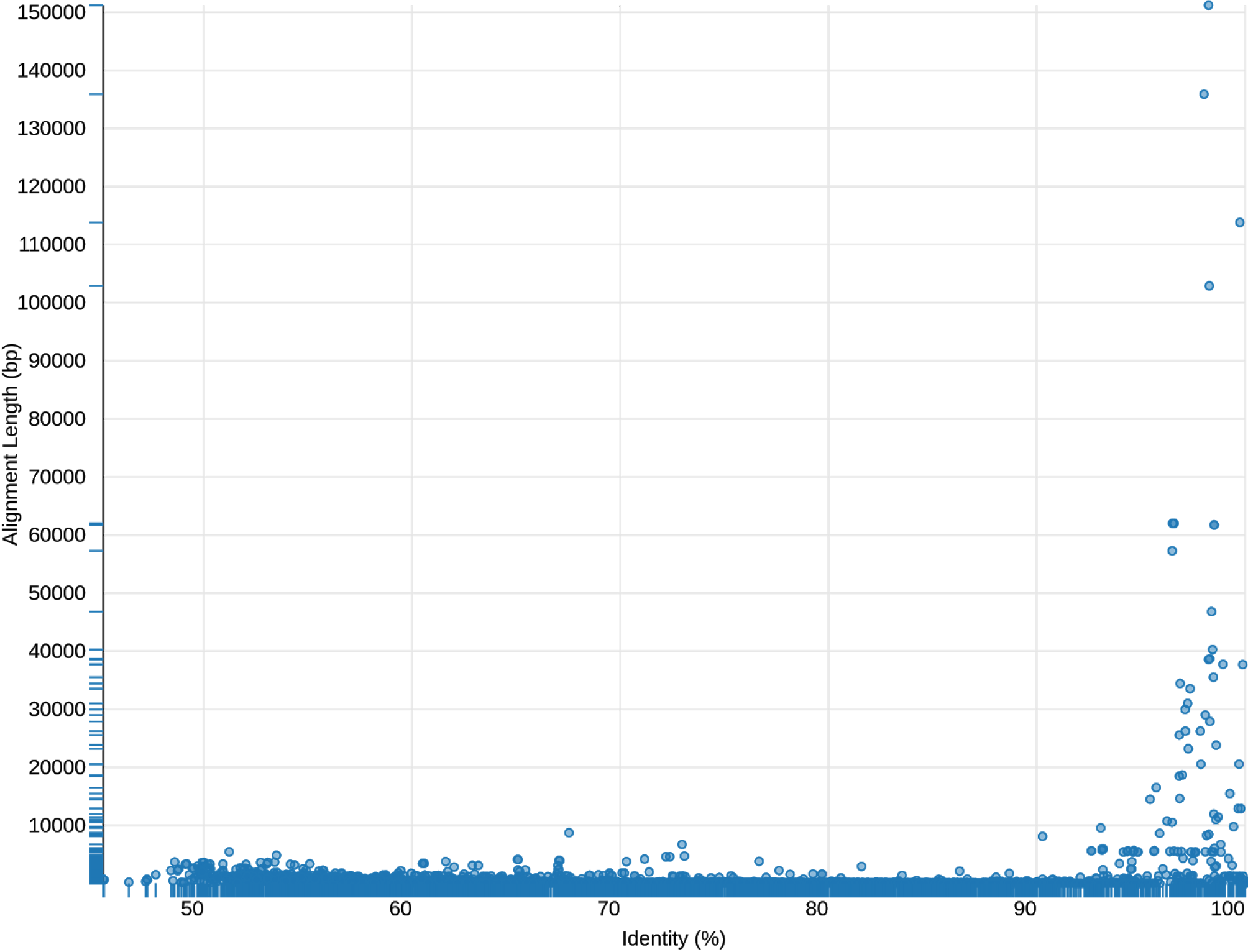
A. The relationship between alignment length and alignment identity allows to identify spurious alignments majority of which are short and have relatively low identity. B. The three genomes most closely related to our assembly appear as a cluster of three dots in the upper left corner of the plot. These have the highest total alignable length over the smallest number of alignment blocks.

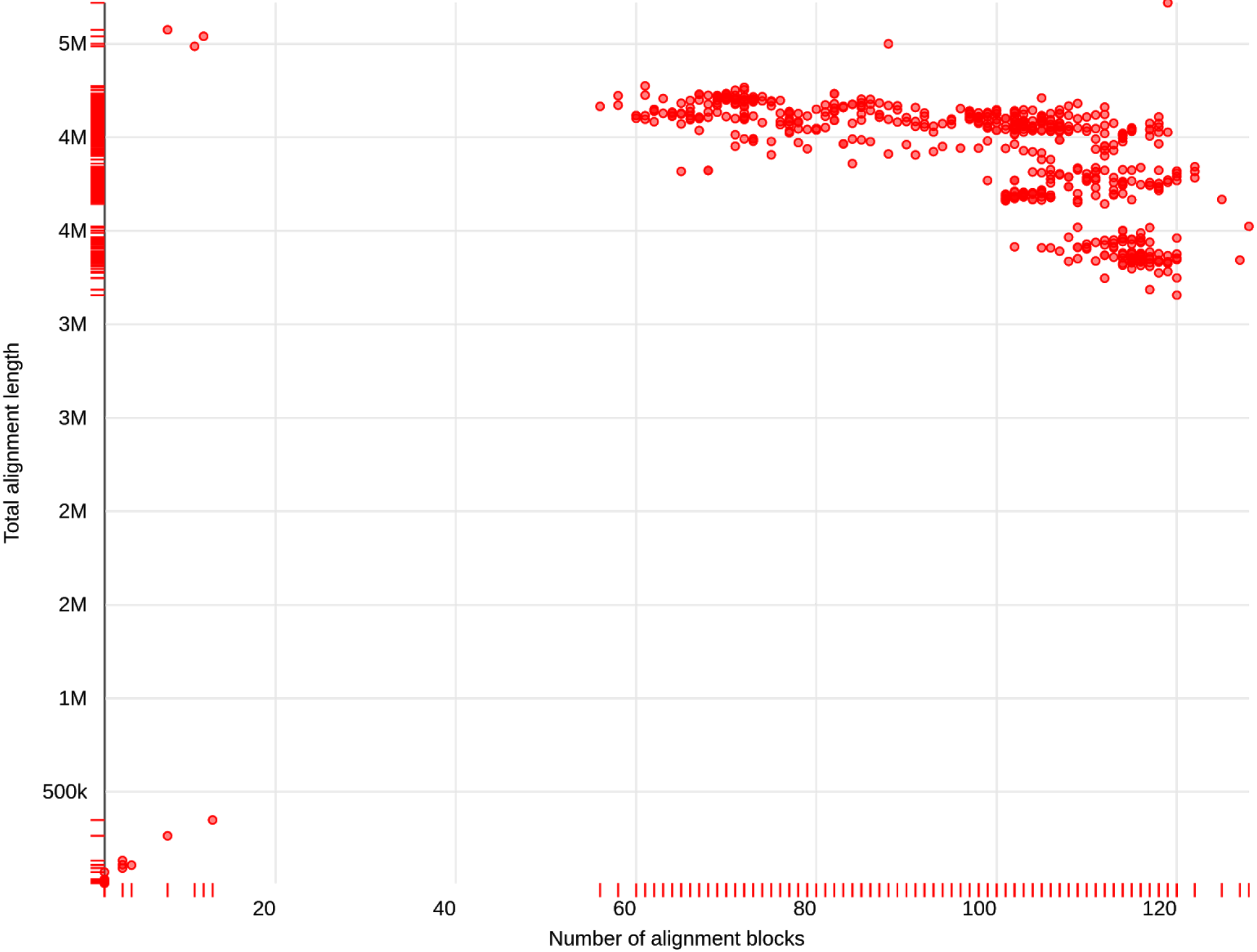

With a set of closely-related genomes identified the next step is to perform a detailed comparison. This can be done by regeneration of alignments. As a bonus LASTZ produces dot-plot representations for alignable regions between every pair of genomes (Fig. 2), highlighting an inversion within CP020543.1 and apparent deletion of approximately 50 kb from our assembly (thus from the three closely related genomes another example of strain C is the most distant from our version of the same strain). It is also possible to create simultaneous representation of pairwise comparisons using tool like Circos ^17^. Circos is a very powerful graphing tool but is also very challenging to configure. To make Circos accessible to wider audience of users we have integrated it into Galaxy to produce a graphical representation of relationships between our assembly and the three genomes (Fig. S1). However, both dot-plots and Circos plots are not interactive. Genome browsers such as the Integrative Genome Viewer (IGV^18^) are designed to allow interactive explorations of genomes and associated annotations. To create a browser one first needs to select a set of sequences that would form the coordinate system for displaying genomic features. In our case we can combine sequences of our assembly with LT906474.1, CP024090.1, and CP020543.1 in a single file. With minor deviations (cleaning sequence names and retaining accession numbers only) this can be accomplished. Once a browser is created, one can display tracks. In the case of our analysis we will display two kinds of tracks: alignment coordinates and gene annotations. Alignment coordinates were produced by LASTZ, and by moving columns around its output can be coerced into Browser Extensible Data (BED) format understood by IGV. The gene annotations can be downloaded from NCBI directly and displayed as tracks. Finally, it is possible to compute a complement of alignment coordinates (essentially an inverse) to pinpoint location of the gap in our assembly (Fig. S2) using BEDTools—a toolkit for manipulation of BED datasets ^19^. However, this last set of manipulation involves a large number of small operations (e.g., removing and re-shuffling columns in tab-delimited files) that are difficult to perform on large datasets outside UNIX environments or Galaxy.

**Figure 2.**
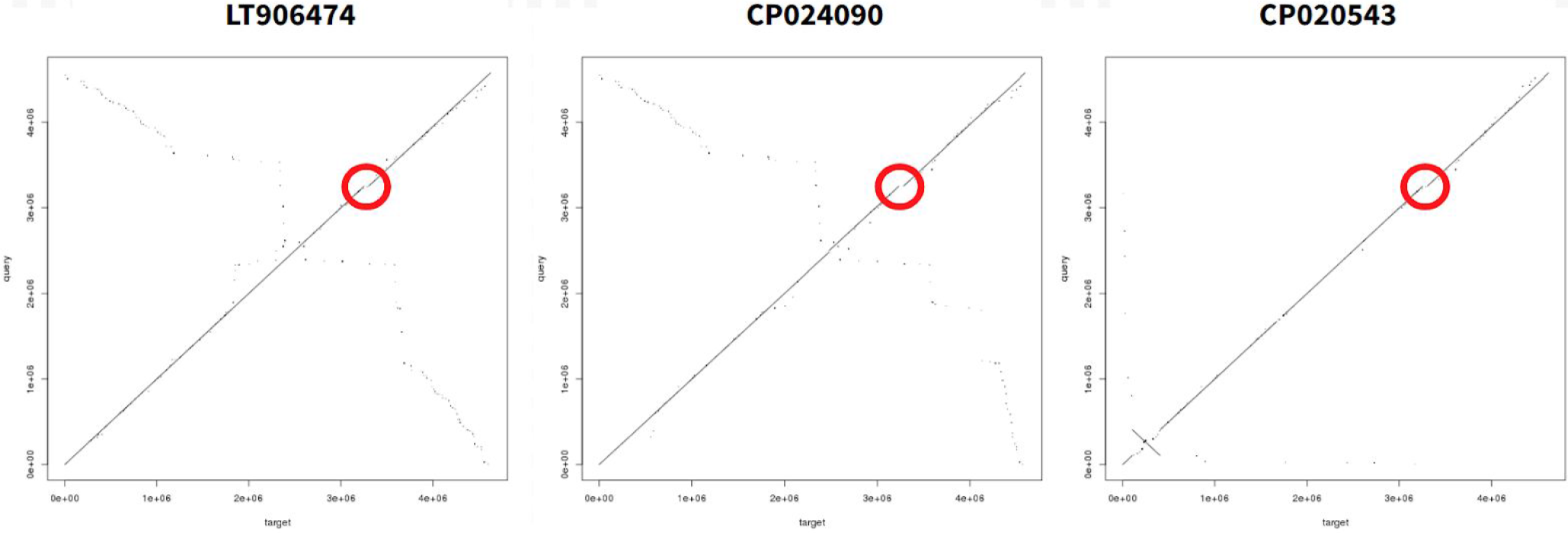
A low resolution comparison of our assembly against the three most closely related genomes identified in Fig. 1B. Query (Y-axis) is indicated above each dot plot. Target (X-axis) is our assembly. Red circle indicates a region deleted in our assembly.

Finally, to understand if any of the genes missing from our assembly (e.g., corresponding to the deleted region in the other three strains) are essential we compared them (using LT906474.1 data) against a recently published list of essential genes identified in *E. coli* K-12 strain BW25113 ^20^. We selected LT906474.1 genes because its genome is better annotated compared with CP020543.1 (e.g., contains standard gene names). The caveat of this analysis is that we are comparing LT906474.1 (WG5) and BW25113—two relatively distant strains (albeit both belonging to the A phylogenetic group of *E. coli*^21^) because there is no systematic gene deletion studies for WG5. This analysis was performed by first identifying genes overlapping the deleted region in LT906474.1 and then comparing their names against those listed by Goodall et al. ^20^. Based on this comparison none of the genes falling within the deleted region appears to be essential.

For experimental co-author of this paper it took approximately two months to figure out numerous minute aspects of the above analysis (computational co-authors deliberately did not provide any help for the duration of this period). After integration of all tools and combining them with existing Galaxy functionality this entire analysis took less than two work-days (because Unicycler assembly and alignment to 592 genomes takes about 12 hours in total). The main take-home-message of our report is that while major analytical problems in bacterial assembly are solved (there are proven, free, open-source, robust tools for assembly, alignment, visualization and so on) the small issues are the true show-stoppers for many experimentalists. These “last-mile” challenges are as mundane as tabular file manipulation, dealing with multiple files, simple scripting and statistical data analysis. Why is this the case and how to overcome these issues?

1. **Lack of quantitative training.** This is likely the most severe problem but also the one that is easiest to solve because of the abundance of free training resources. While we do not necessarily expect biologists to develop quantitative methods for data analytics, sound statistical and scientific computing training is key to successful career. Quantitative courses must become a required component of undergraduate curriculum in all life science programs. This is not a tall order as most universities and large research institutions already have all necessary expertise in one place.
2. **Lack of interdisciplinary crosstalk.** Microbiologists do not often read computational journals. Algorithm developers rarely venture into biological conferences. These sociological trends are very effective in preventing spread of mutually beneficial information across fields. This can potentially change if journals, granting agencies, and domain-specific mindsets would become more receptive to cross-disciplinary efforts or at least acknowledge its importance. Without this, grant proposals describing tailoring of computational tools to the needs of microbiologists will be returned with “no novelty” review statement. And as many of us know “no-novelty” kills it all.

## Acknowledgements

The authors are grateful to Galaxy Team and Galaxy community as without these resources this work would not be possible. This project was supported by NIH Grants U41 HG006620 and R01 AI134384-01 as well as NSF Grant 1661497 to JT and AN.

---

**Supplemental Figure 1.**
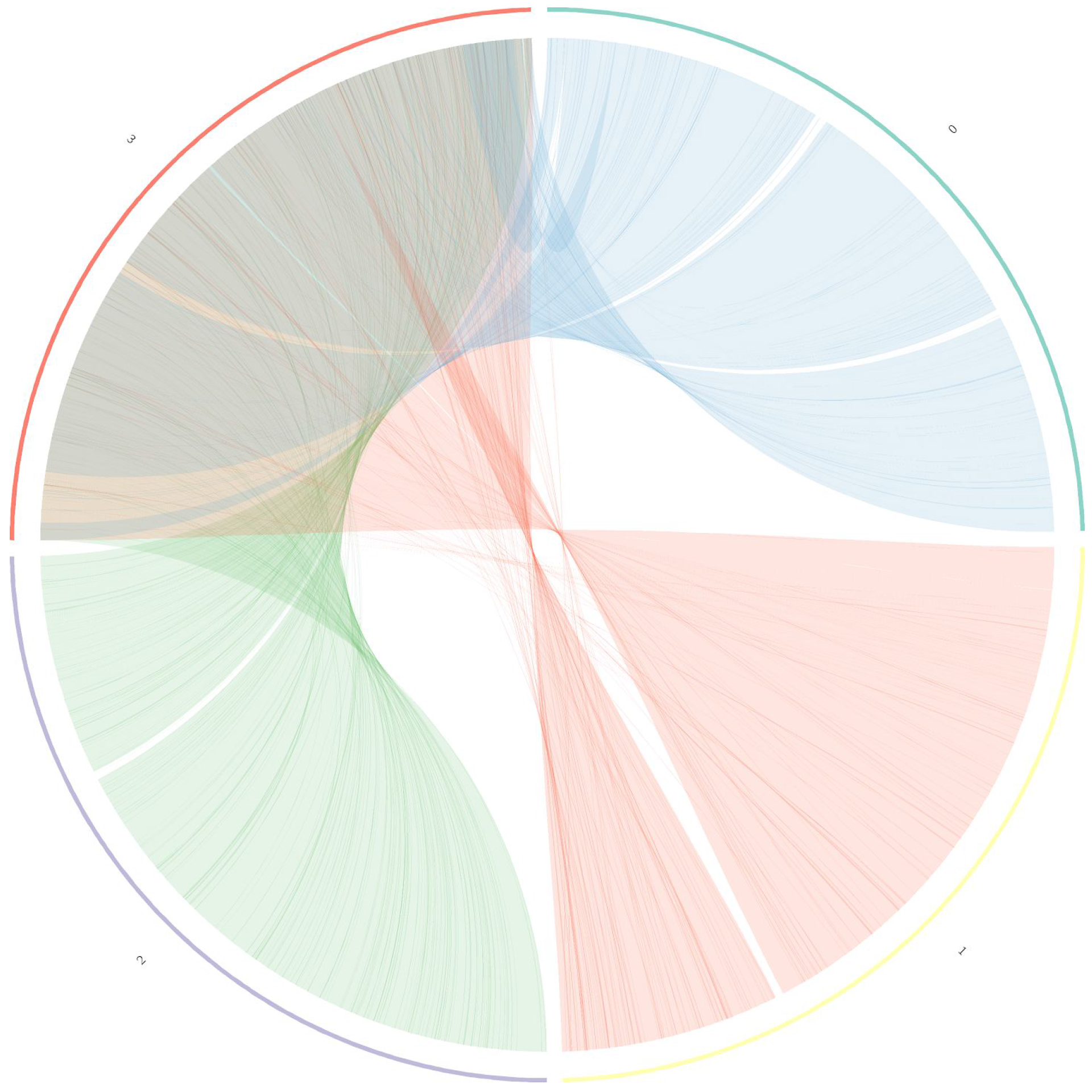
A Circos representation of data shown in Fig. 2. 1 = CP020543.1, 2= CP024090.1, 3 = LT906474.1, 4 = Our assembly

**Supplemental Figure 2.**
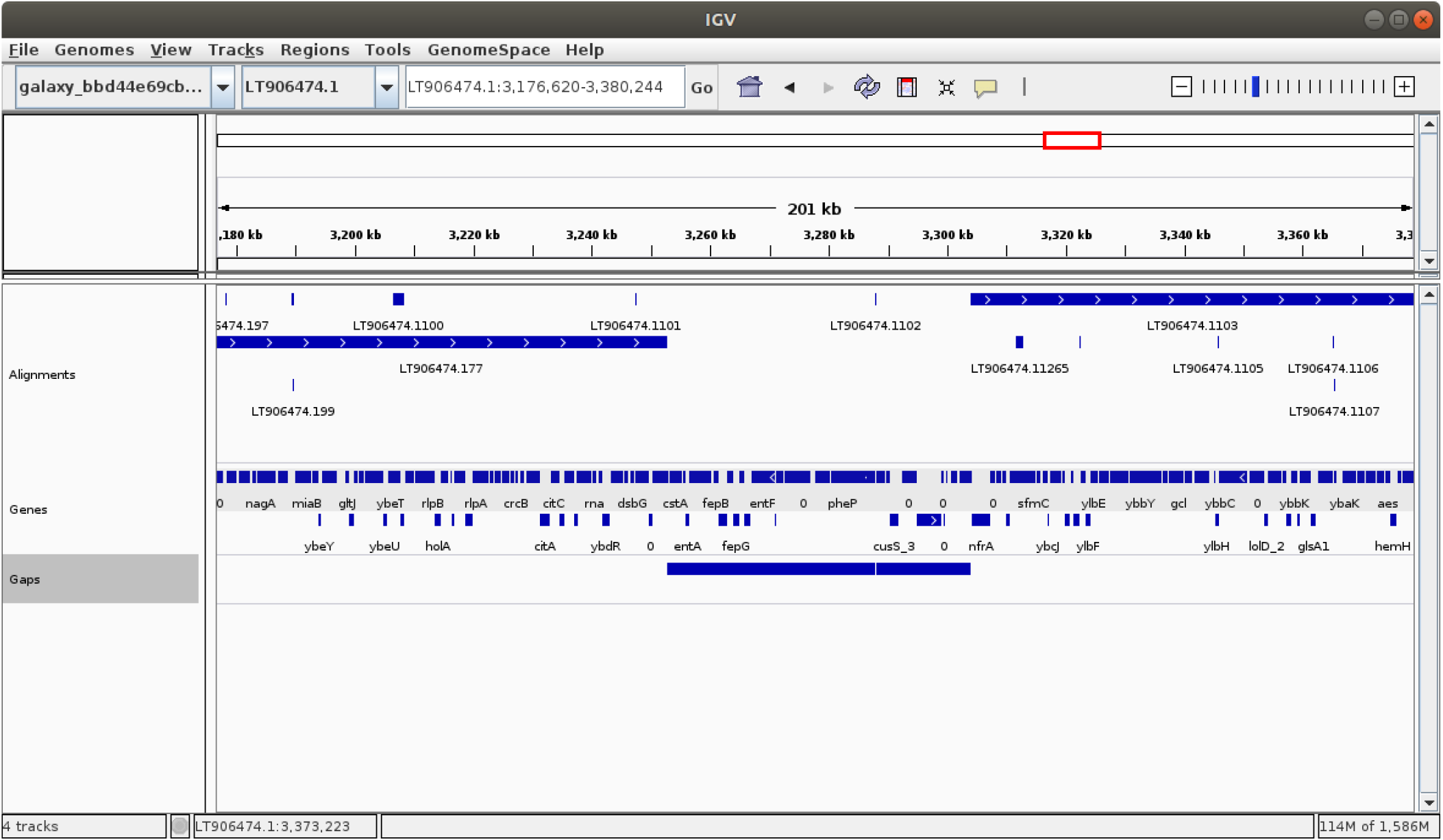
A browser showing alignment, genes, and gaps tracks.

